# Engineered mRNA-packaging virus-like particles enable efficient and precise mitochondrial base editing *in vitro* and *in vivo*

**DOI:** 10.64898/2026.10.04.756548

**Authors:** Hyewon Jang, Carolline Rodrigues Menezes, Krzysztof Palczewski, Aditya Raguram

## Abstract

Mitochondrial DNA (mtDNA) base editors (mtBEs) are promising tools for studying and correcting pathogenic mtDNA mutations. However, the application of mtBEs is often hindered by substantial mtDNA-wide off-target editing that accumulates with prolonged mtBE expression resulting from plasmid DNA or viral delivery. To overcome this limitation, we developed mtBE-VLPs, a virus-like-particle (VLP) platform for transiently delivering mtBEs *in vitro* and *in vivo* with minimal off-target effects. While canonical protein-packaging VLPs yielded undetectable mtDNA base editing, engineered mRNA-packaging VLPs enabled efficient but transient mtBE cargo delivery. By iteratively engineering mRNA-packaging VLP compositions, we improved mtBE-VLP-mediated base editing efficiencies from less than 1% to greater than 50% across multiple target mtDNA loci in bulk populations of treated cultured cells. To further improve the packaging and delivery of certain mtBEs, we developed a strategy to selectively reduce mtBE cargo expression in VLP producer cells but not transduced cells, which increased VLP-mediated base editing efficiencies for certain mtBEs by 7.1-fold. Compared to plasmid DNA delivery *in vitro*, mtBE-VLPs exhibited equivalent on-target editing but 23- to 41-fold lower mtDNA-wide off-target editing, highlighting the improved editing specificity offered by transient mtBE delivery. mtBE-VLPs efficiently installed a pathogenic mtDNA mutation in mouse embryonic fibroblasts (88% efficiency) and corrected a pathogenic mtDNA mutation in a human patient-derived cell line (41% efficiency). Additionally, a single subretinal injection of mtBE-VLPs yielded 6–20% (average 12%) mtDNA base editing in the mouse retinal pigment epithelium with minimal mtDNA-wide off-target editing. These findings establish mRNA-packaging mtBE-VLPs as promising vehicles for efficient and precise mitochondrial base editing via transient mtBE delivery.

---

Mutations in mitochondrial DNA (mtDNA) can disrupt mitochondrial function and contribute to a broad range of human diseases that currently lack effective therapies^1^. The recent development of technologies for mitochondrial genome editing has raised the possibility of using these mtDNA-editing agents to study and potentially treat mitochondrial genetic disorders^2,3^. Specifically, mitochondrial base editors (mtBEs) have enabled targeted C•G-to-T•A and A•T-to-G•C conversions in mtDNA^4,5^, which are sufficient in principle to install and correct 83% of all confirmed pathogenic mtDNA variants^6^. Following the development of the original CRISPR-free mitochondrial cytosine (DdCBE^5^) and adenine (TALED^4,7^) base editors, subsequent engineering has further improved on-target activity^8,9^, expanded the range of targetable sequences^8,10,11^, and enabled strand-specific mtDNA editing^12,13^.

Despite this recent progress, the application of mtBEs as research tools and potential therapeutics has been strongly hindered by off-target editing concerns^2^. While various mtBEs have been used across several cell types and organisms^2,11,14–18^, most studies have detected substantial off-target editing induced by mtBEs, especially DdCBEs. In particular, several reports have demonstrated that adeno-associated-virus (AAV) delivery of mtBEs into mice can cause high levels of mtDNA-wide off-target editing that in some cases can induce severe adverse effects, including premature death^11,14,16^. The prolonged expression of AAV-delivered mtBEs likely contributes to increased off-target editing that accumulates over time in transduced cells^14,19,20^. Therefore, transient delivery of mtBEs could limit off-target effects and provide a more promising approach for mitochondrial base editing *in vitro* and *in vivo*. Indeed, transient delivery of mtBE-encoding mRNAs was shown to reduce off-target effects *in vitro*^21–23^. However, to our knowledge, no strategy for transient post-natal *in vivo* delivery of mtBEs has been previously reported.

Virus-like particles (VLPs) are promising delivery vehicles that have enabled transient delivery of Cas9-based gene editing agents with minimal off-target editing of nuclear DNA *in vitro* and *in vivo*^19^. VLPs use viral scaffolds to package and deliver cargo proteins or RNAs instead of viral genetic material, thereby affording transient cargo delivery rather than prolonged expression from delivered viral transgenes^19^. Additionally, VLPs possess programmable cell-type targeting capabilities and have enabled efficient cargo delivery across a range of cell and tissue types^19,24^. Thus, VLPs could in principle serve as ideal vehicles for transient mtBE delivery, but the compatibility of mtBEs with VLP delivery has thus far been unexplored.

In this study, we report the development of mtBE-VLPs that enable efficient mitochondrial base editing with minimal off-target editing *in vitro* and *in vivo*. Canonical protein-packaging engineered VLPs (eVLPs)^25,26^ yielded undetectable mitochondrial base editing, which led us to identify mRNA delivery instead as a preferred strategy for transient mtBE delivery. By iteratively engineering mRNA-packaging VLP designs, we developed an efficient mRNA-packaging VLP system that improved VLP-mediated mtBE delivery by over 20-fold. The resulting mtBE-VLPs enabled efficient mitochondrial base editing across several mtDNA target sites and cell types *in vitro*, with 23- to 41-fold lower mtDNA-wide off-target editing compared to plasmid delivery. mtBE-VLPs enabled the installation of a disease-relevant mtDNA mutation in mouse embryonic fibroblasts (MEFs), as well as the correction of a pathogenic mtDNA mutation in a patient-derived cell line *in vitro*. Additionally, mtBE-VLPs supported efficient *in vivo* mitochondrial base editing in mouse retinal pigment epithelial (RPE) cells with minimal mtDNA-wide off-target editing. These results establish mtBE-VLPs as a promising platform for efficient and precise mitochondrial base editing.

## Results

### Canonical protein-packaging VLPs do not support mitochondrial base editing

mtBEs such as DdCBEs and TALEDs are large, multi-domain proteins that carry an N-terminal mitochondrial targeting sequence (MTS)^4,8^ and must be imported into the mitochondrial matrix via the TOM/TIM translocase to edit the mitochondrial genome. Such large, multi-domain proteins have been shown to undergo co-translational import into mitochondria prior to folding^27^ (Extended Data Fig. 1a). As a result, if pre-folded MTS-containing proteins are delivered into the target-cell cytosol, their MTSs might not be efficiently recognized by TOM receptors, thereby precluding effective mitochondrial import^28,29^ (Extended Data Fig. 1a). Consistent with this model, we observed that v4 eVLPs^25^ designed to package a pre-folded MTS-containing monomeric DdCBE (mDdCBE)^22^ produced no detectable base editing at the mitochondrial *ND1* target locus across the tested doses (Extended Data Fig. 1b). These findings suggest that mtBE delivery is likely incompatible with canonical protein-packaging gene editing VLPs, necessitating the development of an alternative strategy.

### An optimal component stoichiometry is critical for achieving efficient mRNA-packaging VLP delivery

We reasoned that VLP-mediated delivery of mtBE-encoding mRNAs instead of mtBE proteins could enable co-translational mitochondrial import in target cells and might therefore offer a more promising mtBE delivery strategy (Extended Data Fig. 1a). While protein- and ribonucleoprotein (RNP)-packaging VLPs are most commonly used for delivering gene editing agents, engineered retroviral VLPs have also been used previously to package and deliver mRNA cargos^19,30–33^. In these systems, cargo mRNAs are engineered to contain RNA aptamers that are recognized by specific RNA-binding proteins incorporated into VLP scaffolds (Fig. 1a), which directs the packaging of desired mRNA cargos into VLPs following the co-expression of all components in producer cells. After the transduction of target cells mediated by VLP envelope glycoproteins, the packaged mRNA cargo is delivered into target-cell cytosols, enabling translation of the delivered mRNA cargo and subsequent activity of the translated protein.

**Fig. 1.**
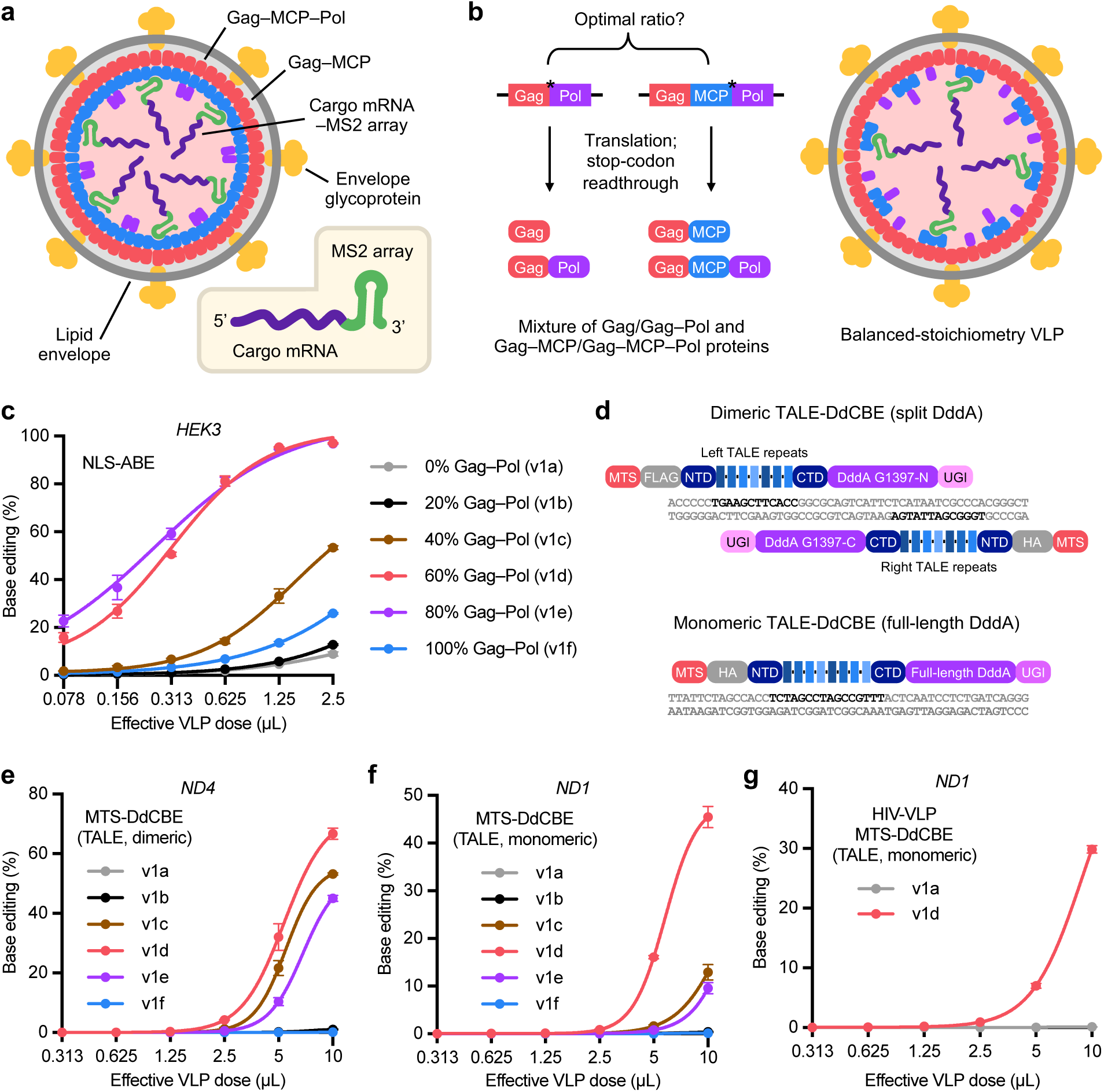
An optimal component stoichiometry is required for functional mRNA-packaging VLPs. **a,** Schematic diagram of the mRNA-packaging VLP platform. MLV Gag–MCP–Pol are co-expressed in producer cells with a VSV-G envelope glycoprotein and a cargo mRNA bearing a 12xMS2 aptamer array in the 3’UTR. The MS2-containing cargo mRNA is selectively packaged into budding, lipid-enveloped particles via the MCP:MS2 interaction. **b,** Schematic diagram of the production of mRNA-packaging VLPs using both WT Gag–Pol and Gag–MCP–Pol. Retroviral stop codon readthrough results in the expression of a mixture of Gag/Gag–Pol and Gag–MCP/Gag–MCP–Pol proteins. * denotes stop codon. **c,** Comparison of nuclear ABE-packaging VLPs produced using different Gag–Pol:Gag–MCP–Pol ratios. Adenine base editing efficiencies at position A_8_ of the nuclear *HEK3* locus in HEK293T cells are shown. **d,** Schematic diagram of the dimeric DdCBE and monomeric DdCBE architectures. Dimeric and monomeric DdCBEs contain split and full-length DddA deaminases, respectively. TALE DNA-binding sites are shown bold. MTS, mitochondrial targeting sequence; TALE, transcription activator-like effector array; NTD, N-terminal domain of split DddA; CTD, C-terminal domain of split DddA; UGI, uracil glycosylase inhibitor. **e,** Comparison of dimeric DdCBE-packaging mtBE-VLPs produced using different Gag–Pol:Gag–MCP–Pol ratios. C•G-to-T•A base editing efficiencies at position C_8_ of the mitochondrial *ND4* locus in HEK293T cells are shown. **f,** Comparison of monomeric DdCBE-packaging mtBE-VLPs produced using different Gag–Pol:Gag–MCP–Pol ratios. C•G-to-T•A base editing efficiencies at position G_10_ of the mitochondrial *ND1* locus in HEK293T cells are shown. **g,** Comparison of monomeric DdCBE-packaging HIV-based VLPs produced using different HIV Gag– Pol:Gag–MCP–Pol ratios. C•G-to-T•A base editing efficiencies at position G_10_ of the mitochondrial *ND1* locus in HEK293T cells are shown. **c, e–g,** Dots and error bars represent mean ± s.d. of n = 3 replicates. Data were fit to four-parameter logistic curves using nonlinear regression. Source data are provided as a Source Data file.

We first began by investigating the ability of mRNA-packaging VLPs to deliver a well-characterized, highly active, nuclear adenine base editor (ABE)^34^. We designed a Murine Leukemia Virus (MLV)-based VLP in which ABE-encoding cargo mRNAs are engineered to contain twelve copies of an MS2 RNA aptamer in the 3’ untranslated region (UTR)^30^ (Fig. 1a, b). MS2 aptamers are recognized by the MS2 coat protein (MCP)^35^, which enables recruitment of MS2-containing cargo mRNAs into VLPs produced using MLV Gag–MCP–Pol scaffolds (Fig. 1a, b). We produced VLPs containing ABE mRNA cargo, MLV Gag–MCP–Pol, and a VSV-G envelope glycoprotein, and we used these VLPs to transduce HEK293T cells that expressed a *HEK3*-targeting guide RNA (gRNA) (see Methods). At the highest VLP doses tested, we detected adenine base editing at the nuclear *HEK3* genomic locus (Fig. 1c). These results confirm and extend previous reports^30,31,33,36^ that the MS2:MCP interaction can enable successful VLP-mediated cargo RNA packaging and delivery *in vitro*.

This initial design of mRNA-packaging VLPs yielded only modest base editing efficiencies, even at high doses of a highly active nuclear base editor. To improve the efficiency of mRNA-packaging VLPs, we next investigated whether the incorporation of wild-type (WT) MLV Gag–Pol in addition to the engineered Gag–MCP–Pol influences delivery efficiencies (Fig. 1b). By varying the amount of WT Gag–Pol-versus Gag–MCP– Pol-encoding plasmids transfected during VLP production (see Methods), we generated a panel of six ABE8e mRNA-packaging VLPs spanning 0–100% WT Gag–Pol abundance in 20% increments (v1a–v1f, Fig. 1c) and evaluated their delivery potency in *HEK3*-gRNA-expressing HEK293T cells, as measured by relative base editing efficiencies. We observed that v1d and v1e VLPs (corresponding to 60% and 80% WT Gag–Pol, respectively) achieved the highest editing efficiencies at the nuclear *HEK3* locus (Fig. 1c), corresponding to a more than 12-fold increase in overall VLP delivery potency (EC_50_). These results indicate that an optimal WT Gag–Pol/Gag–MCP–Pol component stoichiometry is critical for achieving efficient mRNA-packaging VLP delivery.

### Optimized mRNA-packaging VLPs enable efficient DdCBE delivery in vitro

We next evaluated whether these mRNA-packaging VLP compositions could support efficient mtBE delivery and mtDNA base editing. First, we tested VLP-mediated delivery of a dimeric DdCBE with the DddA6 deaminase^8^ targeting the mitochondrial *ND4* locus. Because CRISPR-free mtBEs do not require gRNAs for mtDNA base editing and instead use programmable TALE DNA-binding domains^4,5^ (Fig. 1d), packaging and delivery of mtBE-encoding mRNA is sufficient to edit unmodified target cells. Encouragingly, we observed that dimeric DdCBE-packaging VLPs exhibited efficient cytosine base editing (up to 66% at the highest VLP dose) at the mitochondrial *ND4* locus in HEK293T cells (Fig. 1e). VLPs packaging a monomeric DdCBE^22^ also exhibited efficient cytosine base editing (up to 45% at the highest VLP dose) at the mitochondrial *ND1* locus in HEK293T cells (Fig. 1f). The v1d (60% WT Gag–Pol) composition was similarly optimal for both mtBE-VLP constructs, while the overall delivery potencies varied dramatically across component stoichiometries (Fig. 1e, f), further highlighting the importance of optimizing this parameter for mRNA-packaging VLPs.

To further investigate the effects of different component stoichiometries on mtBE-VLP properties, we quantified mRNA cargo packaging across the different *ND4*-targeting mtBE-VLP constructs using RT-qPCR. We observed that mRNA cargo packaging per particle generally decreased at high abundances of WT Gag–Pol, with v1f (100% WT Gag–Pol) yielding negligible cargo packaging, consistent with progressive loss of the Gag–MCP–Pol required for cargo recruitment (Extended Data Fig. 1c). However, while v1a and v1c VLPs exhibited almost identical cargo packaging (Extended Data Fig. 1c), they exhibited orders-of-magnitude differences in delivery potency (Fig. 1e). Additionally, the most efficient v1d VLPs packaged 3.5-fold fewer cargo mRNAs per particle than the very inefficient v1a VLPs (Extended Data Fig. 1c, Fig. 1e). These results indicate that efficient mRNA cargo packaging is necessary but not sufficient for efficient VLP delivery, and that increased cargo packaging does not necessarily correlate with increased VLP potency. We speculate that insertion of MCP between MLV Gag and Pol might disrupt the tightly regulated Gag:Gag–Pol ratio achieved by retroviral stop codon readthrough^37^, which could in turn impair protease-dependent particle maturation^38^ and subsequent cargo delivery. Therefore, the optimal VLP composition uses the minimum amount of Gag–MCP–Pol required to package sufficient cargo mRNAs and maximizes the amount of WT Gag–Pol incorporated into the particles.

We also explored whether other retroviral scaffolds can support efficient mRNA-packaging mtBE-VLPs. By applying the same MS2/MCP-facilitated cargo packaging strategy, we designed analogous HIV-1-based VLP constructs and evaluated their ability to deliver nuclear ABE or mitochondrial DdCBE mRNAs. We observed that HIV-1-based VLPs also supported efficient ABE and DdCBE delivery into HEK293T cells; the same 60% WT Gag–Pol composition (v1d-HIV) was again more efficient for both constructs (Fig. 1g, Extended Data Fig. 1d). We speculate that the insertion of the MCP domain between HIV Gag and Pol might similarly disrupt Gag:Gag–Pol expression and limit protease-dependent particle maturation^39^. Together, these data indicate that an optimal Gag–Pol:Gag–MCP–Pol stoichiometry is a general requirement for efficient VLP-mediated mRNA delivery across both nuclear and mitochondrial base editor cargos as well as different retroviral scaffolds.

### Reducing envelope glycoprotein expression further improves DdCBE mtBE-VLP potency

After identifying an optimal Gag–Pol:Gag–MCP–Pol stoichiometry, we next investigated whether the stoichiometry of other VLP components—specifically, the envelope glycoprotein VSV-G—influences mtBE-VLP potency. By varying the amount of VSV-G-encoding plasmid transfected during VLP production, we generated six DdCBE mtBE-VLP compositions (v2a–v2f) and evaluated their delivery potencies in mouse Neuro-2a cells (Fig. 2a, b). The optimal v1d Gag–Pol:Gag–MCP–Pol stoichiometry was used in all cases. We observed that reducing the VSV-G plasmid amount generally increased DdCBE mtBE-VLP delivery potency at both the mitochondrial *Nd5* and *Nd4* loci (Fig. 2a, b). The lowest tested VSV-G amount (v2f) led to the greatest improvement in potency, achieving 18-fold and 1.6-fold increases in potency (EC_50_) compared to v2a VLPs at the *Nd5* and *Nd4* loci, respectively. In HEK293T cells, v2f VLPs also improved DdCBE mtBE-VLP potency by 1.8- and 2.4-fold compared to v2a VLPs at the *ND1* and *ATP8* loci, respectively (Fig. 2c, d). However, at the *ND4* locus in HEK293T cells, v2a and v2f DdCBE mtBE-VLPs exhibited similar potencies (Fig. 2e). Additionally, reducing the VSV-G plasmid amount had the opposite effect on ABE-VLP potency: v2b–v2f ABE-VLPs exhibited reduced editing efficiencies compared to v2a ABE-VLPs (Extended Data Fig. 2a). These results suggest that different VLP cargos require distinct optimal VSV-G expression levels, possibly due to cargo-dependent differences in the total protein expression burden in producer cells during VLP production.

**Fig. 2.**
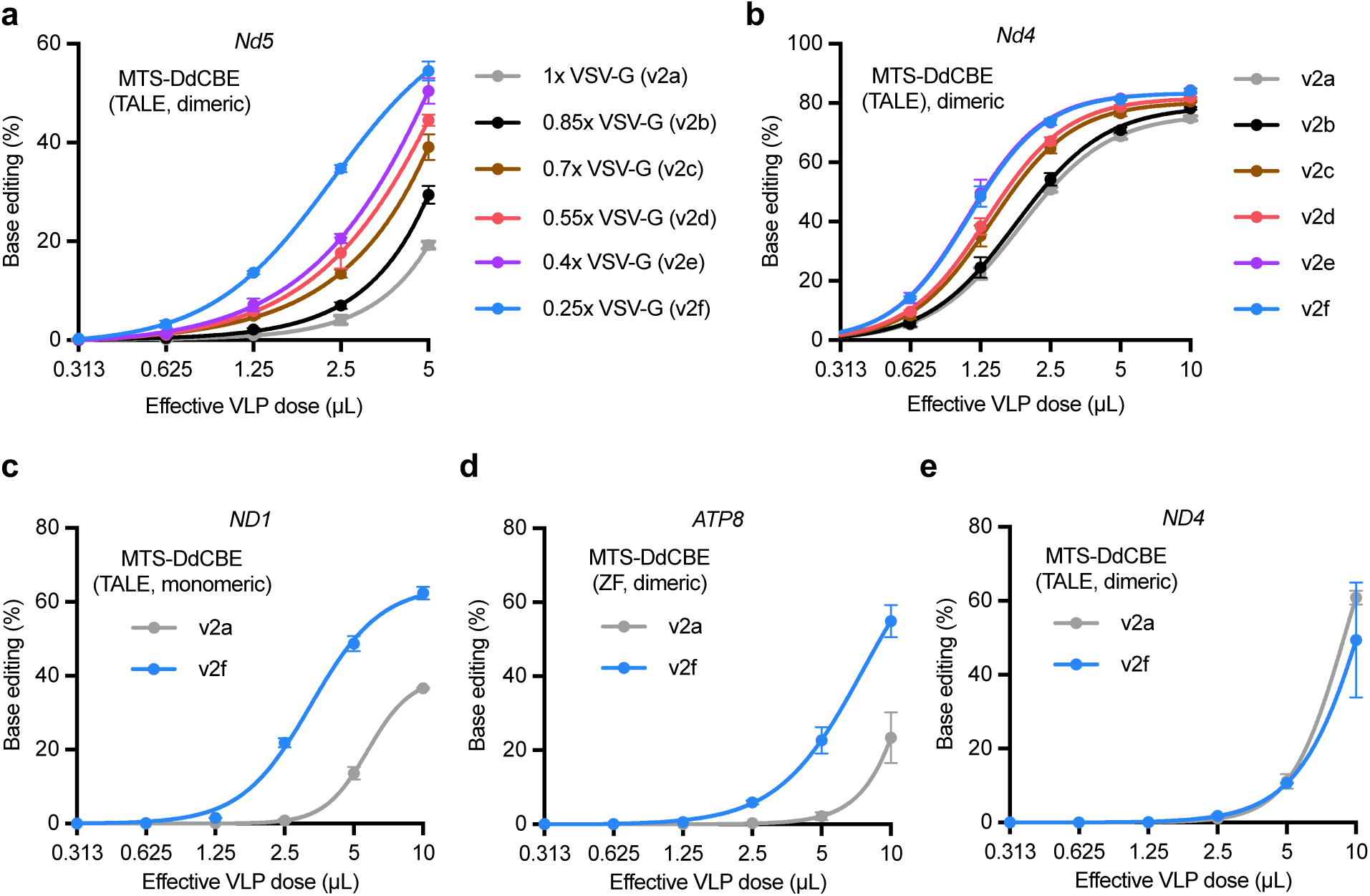
Optimizing envelope glycoprotein expression further improves DdCBE mtBE-VLP potency. **a–b,** Comparison of DdCBE mtBE-VLPs produced using different VSV-G plasmid amounts. C•G-to-T•A base editing efficiencies in mouse Neuro-2a cells are shown at position G_7_ of the mitochondrial *Nd5* locus (**a**) and position G_9_ of the mitochondrial *Nd4* locus (**b**). **c–e,** Comparison of DdCBE mtBE-VLPs at additional mtDNA target sites in HEK293T cells. C•G-to-T•A base editing efficiencies are shown at position G_10_ of the *ND1* locus (**c**), position C_6_ of the *ATP8* locus (**d**), and position C_8_ of the *ND4* locus (**e**). **a–e,** Dots and error bars represent mean ± s.d. of n = 3 replicates. Data were fit to four-parameter logistic curves using nonlinear regression. Source data are provided as a Source Data file.

To further characterize the effect of reducing VSV-G expression, we quantified mRNA cargo packaging in DdCBE mtBE-VLPs. We observed that v2f DdCBE mtBE-VLPs packaged 2.3–4.2-fold more cargo mRNA per particle than v2a DdCBE mtBE-VLPs, consistent with their improved potency (Extended Data Fig. 2b). Since v2f DdCBE mtBE-VLPs always offered equivalent or greater efficiencies compared to v2a DdCBE mtBE-VLPs (Fig. 2a–e), we selected v2f as the final composition used for all subsequent DdCBE mtBE-VLPs. Collectively, these findings reveal that VSV-G expression is an additional, independently tunable parameter for maximizing mRNA-packaging mtBE-VLP potency.

### DdCBE mtBE-VLP delivery substantially reduces mtDNA-wide off-target editing compared to plasmid DNA transfection in vitro

Given that v2f DdCBE mtBE-VLPs enabled robust on-target editing across several mtDNA target sites, we next evaluated mtDNA-wide off-target editing generated by these mtBE-VLPs. We delivered the same DdCBE constructs into HEK293T cells via v2f mtBE-VLPs or plasmid DNA transfection and assessed mtDNA-wide off-target editing at three days post-delivery by whole mitochondrial genome sequencing^7^ (see Methods). Notably, we observed that DdCBE mtBE-VLPs generated substantially fewer off-target C•G-to-T•A mutations distributed throughout the mitochondrial genome (Fig. 3a–c, Supplementary Fig. 1) while achieving comparable or greater on-target editing compared to both low- (30 ng) and high- (100 ng) dose plasmid transfection (Fig. 3d). Specifically, *ND1*-targeting mtBE-VLPs exhibited average mtDNA-wide off-target editing levels that were 5.4- and 41-fold lower than those of low- or high-dose plasmid transfection, respectively; *ND4*-targeting mtBE-VLPs displayed average mtDNA-wide off-target editing levels that were 3.5- and 23-fold lower than those of low- or high-dose plasmid transfection, respectively. Furthermore, *ND1*- and *ND4*-targeting mtBE-VLPs exhibited 13- and 13-fold improvements in on-target:off-target editing ratios compared to low-dose plasmid transfection, as well as 32- and 24-fold improvements compared to high-dose plasmid transfection (Fig. 3e). Interestingly, across both delivery methods, dimeric *ND4*-targeting DdCBEs exhibited lower mtDNA-wide off-target editing compared to monomeric *ND1*-targeting DdCBEs (Fig. 3d, e), likely due to the non-split DddA deaminase domain present in the monomeric *ND1*-targeting DdCBE^22^ that is more prone to off-target deamination. Together, these results demonstrate that mtBE-VLPs exhibit robust on-target editing but substantially lower mtDNA-wide off-target editing compared to plasmid delivery *in vitro*, highlighting the benefits of transient mRNA delivery of mtBEs.

**Fig. 3.**
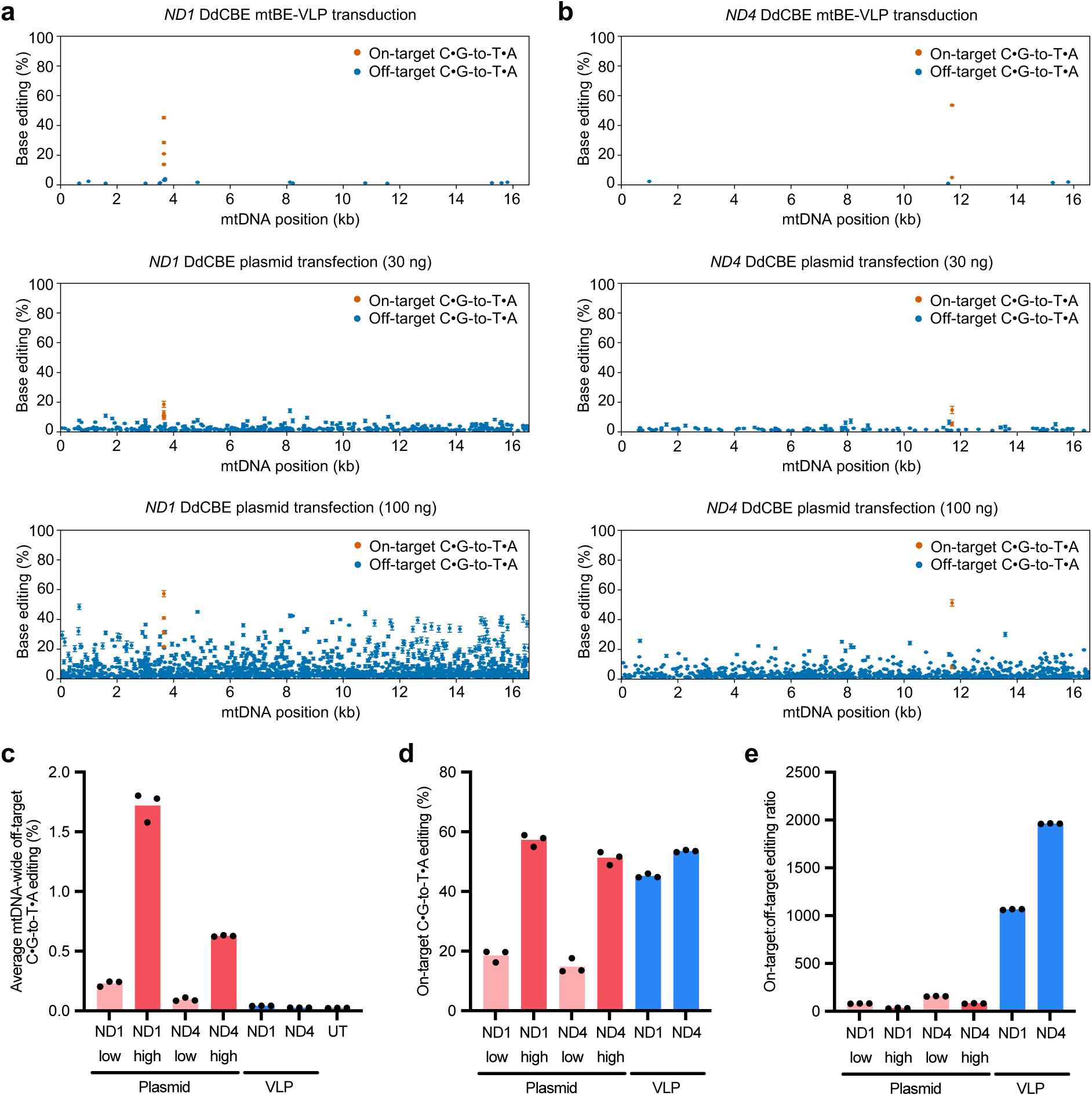
mRNA-packaging mtBE-VLP delivery of DdCBE substantially reduces mtDNA-wide off-target editing compared to plasmid DNA transfection. **a,** mtDNA-wide single-base C•G-to-T•A editing frequencies in HEK293T cells following *ND1*-targeting v2f DdCBE mtBE-VLP transduction or *ND1*-targeting DdCBE plasmid DNA transfection. **b,** mtDNA-wide single-base C•G-to-T•A editing frequencies in HEK293T cells following *ND4*-targeting v2f DdCBE mtBE-VLP transduction or *ND4*-targeting DdCBE plasmid DNA transfection. **a–b,** Each dot represents a single C•G base pair, and error bars represent mean ± s.d. of n = 3 replicates. Orange dots denote on-target editing; blue dots, off-target editing. **c,** Average mtDNA-wide off-target C•G-to-T•A editing frequencies generated by mtBE-VLP transduction or plasmid transfection. UT, untreated. **d**, On-target mtDNA C•G-to-T•A editing efficiencies at the intended *ND1* (position G_10_) or *ND4* (position C_8_) loci generated by mtBE-VLP transduction or plasmid transfection. **e,** On-target:off-target editing ratios associated with mtBE-VLP transduction or plasmid transfection. **c–e,** Bars reflect the mean of n = 3 replicates, and dots represent individual replicate values. Source data are provided as a Source Data file.

### Selective reduction of TALED expression in producer cells enables TALED mtBE-VLP delivery

We next sought to extend the mtBE-VLP platform to package and deliver TALED cargos in addition to DdCBEs. TALEDs enable A•T-to-G•C conversions in mtDNA by combining a TadA8e deaminase with a catalytically inactive DddA and a DNA-binding TALE array^4,7^. We first screened TadA8e variants (WT, V28R, V106W, R111S) for their ability to support TALED mtBE-VLP-mediated editing at the *ND1* locus in HEK293T cells.

In this context, we observed that TadA8e^V106W^ was the most efficient deaminase variant (Fig. 4a), so we used this deaminase variant in all subsequent TALED constructs. However, these TALED mtBE-VLPs only achieved 4% editing at the *ND1* locus at the highest VLP dose tested (Fig. 4a), which was much lower than what was achieved with DdCBE mtBE-VLPs (Fig. 2c), suggesting that additional optimization of TALED mtBE-VLPs was required.

**Fig. 4.**
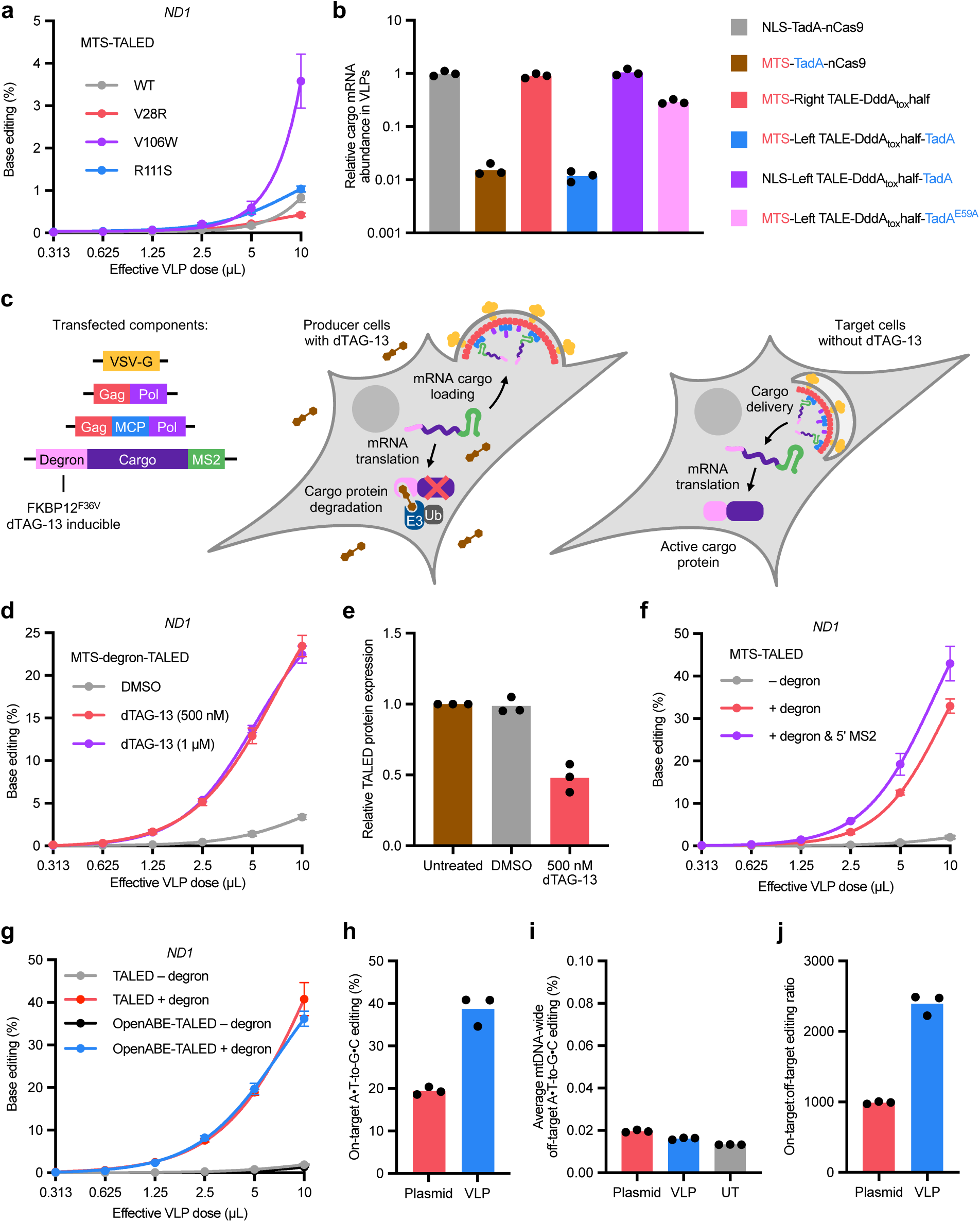
Selective reduction of TALED expression in producer cells enables TALED mtBE-VLP delivery. **a,** Comparison of TALED mtBE-VLPs containing different TadA8e variants. A•T-to-G•C base editing efficiencies at position A_11_ of the mitochondrial *ND1* locus in HEK293T cells are shown. **b,** Relative cargo mRNA abundance across VLP constructs. **c,** Schematic diagram of the degron-based producer-cell expression control system. TALED is fused to an MTS and an FKBP12^F36V^ degron tag. Addition of the small molecule dTAG-13 to producer-cell media induces E3 ligase-mediated degradation of degron-tagged TALED, limiting its expression during VLP production but retaining its expression in transduced cells that are not cultured with dTAG-13. Ub, ubiquitin. **d,** Comparison of degron-containing TALED mtBE-VLPs produced in the presence or absence of dTAG-13. A•T-to-G•C base editing efficiencies at position A_11_ of the mitochondrial *ND1* locus in HEK293T cells are shown. **e,** Relative TALED protein expression in producer cells, normalized to expression in untreated cells. **f,** Comparison of TALED mtBE-VLPs containing different deaminase variants, with or without a degron tag, and in the presence or absence (DMSO) of dTAG-13 during production. A•T-to-G•C base editing efficiencies at position A_11_ of the mitochondrial *ND1* locus in HEK293T cells are shown. **g,** Comparison of degron- and 5’ MS2-containing TALED mtBE-VLPs. All degron-containing mtBE-VLPs were produced in the presence of 500 nM dTAG-13. A•T- to-G•C base editing efficiencies at position A_11_ of the mitochondrial *ND1* locus in HEK293T cells are shown. **h,** On-target mtDNA A•T-to-G•C editing efficiencies at the intended *ND1* (position A_11_) locus in HEK293T cells generated by mtBE-VLP transduction or plasmid transfection. **i,** Average mtDNA-wide off-target A•T-to-G•C editing frequencies in HEK293T cells generated by mtBE-VLP transduction or plasmid transfection. UT, untreated. **j,** On-target:off-target editing ratios associated with mtBE-VLP transduction or plasmid transfection. **a, d, f–g,** Dots and error bars represent mean ± s.d. of n = 3 replicates. Data were fit to four-parameter logistic curves using nonlinear regression. **b, e, h–j,** Bars reflect the mean of n = 3 replicates, and dots represent individual replicate values. Source data are provided as a Source Data file.

To further investigate the properties of TALED mtBE-VLPs, we quantified VLP cargo packaging across several mRNA cargos. We observed that mRNA cargo encoding the MTS–TadA-containing TALED half packaged approximately 100-fold less efficiently than cargo encoding the TadA-free MTS–TALED half (Fig. 4b), suggesting that the expression of MTS–TadA in producer cells was contributing to decreased VLP packaging. Additionally, replacing the nuclear localization signal (NLS) in a TadA-containing nuclear base editor with an MTS similarly resulted in decreased packaging, whereas replacing the MTS of TALED with an NLS restored packaging (Fig. 4b). These results indicate that the VLP packaging defect arises specifically from expressing an MTS–TadA fusion in producer cells rather than expressing either domain alone. Notably, a catalytically inactive MTS–TadA^E59A^ TALED mutant was more efficiently packaged compared to the canonical MTS–TadA TALED (Fig. 4b), suggesting that mitochondrial-localized adenosine deaminase activity in producer cells could be the primary driver of the observed packaging defect.

We reasoned that selectively reducing TALED protein expression during packaging in producer cells, while still allowing VLP-delivered TALED mRNA to be translated in target cells, could help alleviate the TALED packaging defect. To achieve this selective expression, we incorporated a degron tag (FKBP12^F36V^) near the N-terminus of TALED that triggers targeted protein degradation in the presence of a small-molecule inducer, dTAG-13^40,41^ (Fig. 4c). Therefore, in producer cells cultured in dTAG-13-containing media, TALED protein expression should be reduced, without affecting TALED mRNA expression (the latter of which is required for VLP cargo packaging). However, in target cells cultured in media without dTAG-13, TALED proteins expressed from the delivered mRNA cargo should be maintained without degradation (Fig. 4c). Encouragingly, we observed that this degron-addition strategy increased maximal TALED mtBE-VLP efficiencies at the *ND1* locus in HEK293T cells by 7.1-fold relative to DMSO or untreated controls (Fig. 4d, Supplementary Fig. 2a). Incorporation of the degron into the cargo construct and the addition of ≥ 500 nM dTAG-13 to the producer-cell culture media were both required to achieve this improvement in TALED mtBE-VLP potency (Fig. 4d, Supplementary Fig. 2b). Additionally, we confirmed via Western blot that TALED protein levels in producer cells transfected with an HA-tagged degron-containing cargo plasmid were 2.1-fold lower when dTAG-13 was added (Fig. 4e, Supplementary Fig. 2c). These data indicate that this inducible-degron strategy can successfully diminish protein expression in producer cells and can improve the potency of mRNA-packaging TALED mtBE-VLPs.

Based on these findings, we reasoned that TALED’s deaminase activity in producer-cell mitochondria was negatively impacting VLP production in some way. Mitochondrial membrane potential and cellular viability levels were not substantially affected by TALED expression (Supplementary Fig. 3). However, we observed that TALED expression resulted in substantially elevated A-to-I off-target editing of producer-cell mitochondrial RNA relative to DdCBE, NLS-ABE, TALED^E59A^, and untreated controls (Supplementary Fig. 4), consistent with previous reports of TALED’s extensive cellular transcriptome-wide off-target RNA editing^7^. Producer-cell TALED degradation using the degron strategy resulted in a partial reduction in A-to-I off-target editing of mitochondrial RNA (Supplementary Fig. 4). These data suggest that increased TALED-induced off-target editing of mitochondrial RNA in producer cells correlates with decreased VLP packaging and delivery potency, and that this off-target RNA editing can be partially mitigated by the degron strategy described above. While the detailed mechanism of TALED-mediated packaging impairment warrants further investigation, our findings nonetheless indicate that the inducible-degron strategy enables efficient TALED mtBE-VLP production and could offer a general way to deplete problematic producer-cell cargo protein expression that might be relevant for other mRNA-packaging VLPs.

Lastly, we further characterized the properties of TALED mtBE-VLPs. Increasing the number of dTAG-13 treatments in producer cells or titrating the VSV-G amount did not further improve delivery potency (Supplementary Fig. 2d, e), whereas incorporating an MS2 aptamer into the 5’ UTR modestly improved delivery potency (Fig. 4f). Furthermore, we found that the inducible-degron strategy was essential for achieving improved mtDNA editing efficiencies at the *Nd4* and *Nd5* loci in Neuro-2a cells (Extended Data Fig. 3). mtBE-VLPs also enabled the delivery of OpenABE-TALED^42^ cargo, yielding editing efficiencies comparable to those of standard TadA-TALED at the *ND1* locus in HEK293T cells (Fig. 4g). Additionally, TALED mtBE-VLPs achieved 2-fold greater on-target editing compared to plasmid delivery at the *ND1* locus in HEK293T cells (Fig. 4h); both TALED mtBE-VLPs and TALED plasmid delivery exhibited minimal mtDNA-wide off-target editing (Fig. 4i, j, Extended Data Fig. 4), consistent with the improved DNA specificity of TALEDs compared to DdCBEs^7^. Taken together, our results extend the mtBE-VLP platform to both DdCBEs and TALEDs, two important classes of mtDNA base editors.

### mtBE-VLPs mediate robust installation and correction of pathogenic mtDNA mutations in vitro

We next sought to apply mtBE-VLPs to install and correct pathogenic mtDNA mutations in different cell types. The most common primary mtDNA disorder is Leber hereditary optic neuropathy (LHON), which leads to acute or subacute blindness in young adults and affects roughly one in 30,000 individuals^43^. The *ND4* m.G11778A (R340H) mutation, one of three mtDNA point mutations that cause the vast majority of LHON cases, is found in 50–70% of patients of European ancestry and in over 90% of patients with East Asian ancestry^44,45^. First, we used DdCBE mtBE-VLPs to install the mouse *Nd4* m.G11185A mutation, which mimics the human pathogenic *ND4* m.G11778A mutation^15^, in primary mouse embryonic fibroblasts (MEFs). We observed that v2f DdCBE mtBE-VLPs achieved up to 88% installation of the target mutation at the mouse mitochondrial *Nd4* locus (Fig. 5a). Since the generation of homoplasmic mitochondrial disease models requires high on-target editing efficiency, these results suggest that mtBE-VLPs could be valuable for the future generation of mouse models of mitochondrial disease. mtBE-VLPs could be especially compatible with a recently described approach that generated genetically engineered mouse models using simple co-incubation of zygotes and nuclear gene editing VLPs, without the need for a complex microinjection setup^46^.

**Fig. 5.**
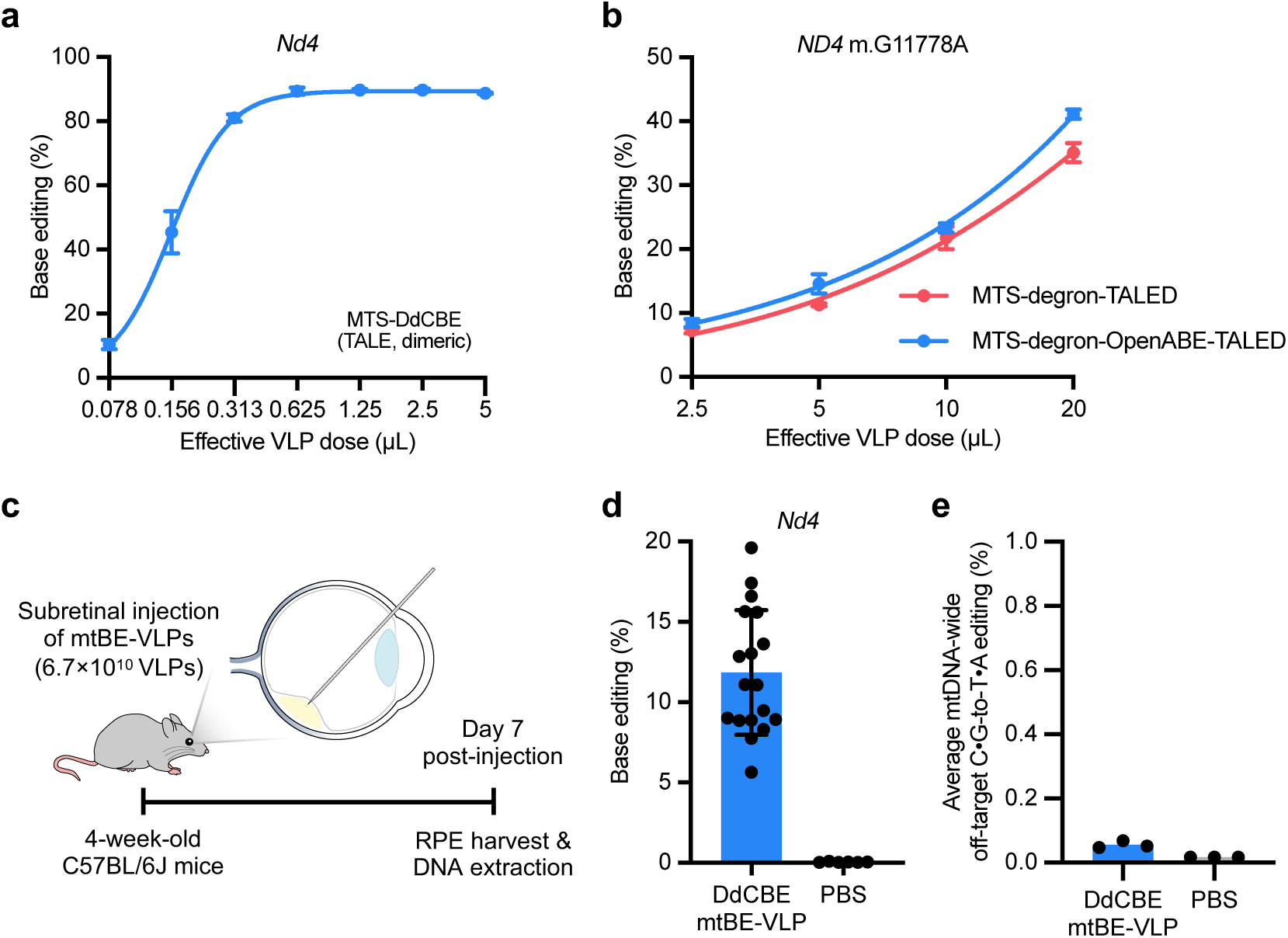
mtBE-VLPs mediate robust mitochondrial base editing in primary cells and *in vivo*. **a,** Efficiency of installing the LHON-disease-relevant m.G11185A mutation (C•G-to-T•A editing at position G_9_) at the mitochondrial *Nd4* locus in primary MEFs mediated by DdCBE mtBE-VLPs. **b,** Comparison of degron- and 5’ MS2-containing TALED mtBE-VLPs in LHON-patient-derived LCLs. All VLPs were produced in the presence of 500 nM dTAG-13. Correction efficiencies of the *ND4* m.G11778A mutation, corresponding to A•T-to-G•C base editing at position A_2_, are shown. **c,** Schematic diagram of the *in vivo* subretinal injection study design. **d,** On-target C•G-to-T•A editing at position G_9_ of the mitochondrial *Nd4* locus in bulk retinal pigment epithelium (RPE) tissue isolated from DdCBE mtBE-VLP-injected or PBS-injected eyes. Bars reflect the mean ± s.d. of n = 18 mice (VLP injection) and n = 6 mice (PBS injection), and dots represent individual mouse values. **e,** Average mtDNA-wide off-target C•G-to-T•A editing frequencies in bulk RPE isolated from DdCBE mtBE-VLP-injected or PBS-injected eyes. Bars reflect the mean of n = 3 mice, and dots represent individual replicate values. **a–b,** Dots and error bars represent mean ± s.d. of n = 3 replicates. Data were fit to four-parameter logistic curves using nonlinear regression. Source data are provided as a Source Data file.

We next evaluated whether TALED mtBE-VLPs could correct a LHON-disease-causing mutation in a relevant human-cell context. We transduced a LHON patient-derived lymphoblastoid cell line (LCL) harboring the *ND4* m.G11778A mutation with mtBE-VLPs packaging either standard TadA-TALED or OpenABE-TALED mtBEs. These mtBE-VLPs achieved up to 41% correction of the pathogenic mutation at the highest VLP dose tested (Fig. 5b). This correction efficiency exceeds by 2-fold the ∼20% efficiency achieved in the same cell line in a previous study that used electroporation of a circular RNA encoding a mitochondrial adenine base editor^13^. Collectively, these results demonstrate robust mtBE-VLP-mediated installation and correction of pathogenic mtDNA mutations in relevant cell types, highlighting the utility of mtBE-VLPs for disease modeling and potential mtDNA-editing therapies.

### mtBE-VLPs enable efficient in vivo mitochondrial base editing with minimal mtDNA-wide off-target editing

Finally, we applied mtBE-VLPs to achieve *in vivo* mitochondrial base editing in the mouse retinal pigment epithelium (RPE). Several recent reports have demonstrated efficient VLP-mediated nuclear gene editing in the mouse RPE via subretinal injection of VLPs^25,36,47–49^. We sought to evaluate whether mtBE-VLPs could install the LHON-disease-relevant *Nd4* m.G11185A mutation in wild-type mouse RPE as a proof-of-concept demonstration of *in vivo* mtDNA base editing. Subretinal injection of v2f DdCBE mtBE-VLPs (6.7×10^10^ particles) into wild-type adult mice yielded 6–20% (average 12%) C•G-to-T•A base editing at the *Nd4* target locus in bulk RPE tissue isolated 7 days post injection (Fig. 5c, d). This mtDNA editing efficiency is comparable to nuclear DNA editing efficiencies in the mouse RPE that were previously achieved by nuclear gene editing VLPs^25,36,47–49^; however, given the difference in the per-cell copy number of mtDNA (typically thousands^50^) versus nuclear DNA (generally two), mtBE-VLPs must edit many more DNA substrates compared to nuclear gene editing VLPs to achieve high bulk editing efficiencies.

We next performed whole mitochondrial genome sequencing to evaluate *in vivo* mtDNA-wide off-target editing. In DNA isolated from three mtBE-VLP-treated RPE samples that exhibited the highest on-target editing, we observed minimal (average 0.06%) mtDNA-wide off-target editing (Fig. 5e, Supplementary Fig. 5). Notably, our results contrast with previous studies that used AAV delivery of DdCBEs in mice and detected substantially higher mtDNA off-target editing (specifically, 0.22–0.30% average mtDNA-wide off-target editing^16^ or many off-target cytosines edited at greater than 5% efficiency^11,14^), likely due to the accumulation of off-target editing over weeks of AAV-delivered mtBE expression^20^. These results demonstrate that transient mtBE delivery mediated by mtBE-VLPs can achieve robust *in vivo* mitochondrial base editing with minimal mtDNA-wide off-target editing, establishing mtBE-VLPs as a promising alternative to existing viral delivery methods for *in vivo* mtBE delivery.

## Discussion

In this study, we developed mRNA-packaging mtBE-VLPs that enable efficient, transient delivery of mtBEs *in vitro* and *in vivo*. By identifying VLP component stoichiometry as a previously underappreciated parameter that strongly influences mRNA-packaging VLP potency, we improved the delivery efficiency of mtBE-VLPs by over 20-fold through iterative engineering. mtBE-VLPs enabled efficient correction of a pathogenic mtDNA mutation in a patient-derived cell line and *in vivo* mitochondrial base editing with minimal off-target editing in the mouse RPE.

The development of optimized mRNA-packaging VLP compositions was essential for enabling efficient mtBE cargo packaging and delivery. While protein- and RNP-packaging VLPs have been most commonly used for delivering Cas9-based gene editing agents^19^, we found that these canonical VLPs yielded undetectable mtDNA base editing, likely due to the distinct requirements for efficient nuclear versus mitochondrial import of proteins in target cells. mRNA delivery, which enables co-translational mitochondrial import of MTS-containing mtBEs in target cells, was required for efficient VLP-mediated mtDNA base editing.

Some mRNA-packaging VLPs have been previously used to package and deliver Cas9 mRNAs^31,33,51^, but challenges associated with separate guide RNA-packaging requirements have complicated their use. Because mtBEs do not require guide RNAs, they are particularly suitable cargos for mRNA-packaging VLPs. Additionally, unlike protein packaging, mRNA packaging in principle allows for the selective expression of cargos in transduced cells but not in producer cells; we achieved this selectivity in practice by incorporating small-molecule-inducible degron tags into mtBE cargos, which enabled selective degradation in producer cells of specific mtBEs whose expression appeared to disrupt VLP production. We anticipate that this selective cargo expression strategy could also be useful for other cargos beyond specific mtBEs. In addition to cargo engineering, producer-cell engineering^52^ could provide a complementary strategy for further improving mtBE-VLP cargo packaging and production.

Our results demonstrate that mtBE-VLPs provide a general platform for transient mtBE delivery *in vitro* and *in vivo*, which is critical for minimizing deleterious mtDNA-wide off-target editing. While previous studies have explored transient delivery of mtBEs *in vitro*^21–23^, the application of transient mtBE delivery *in vivo* demonstrated in our study strongly improves the therapeutic potential of mtBEs. In particular, the use of mtBE-VLPs could minimize the off-target editing and related toxicity concerns associated with AAV-mediated mtBE delivery^11,14^, which is the only method that has previously been used for *in vivo* mtBE delivery, to our knowledge. mRNA-packaging lipid nanoparticles (LNPs), which have shown promise for transient delivery of nuclear gene editing agents^19,53^, could in principle also be used to achieve transient *in vivo* mtBE delivery, although this has not been demonstrated to date. Compared to LNPs, VLPs offer several advantages, including programmable cell-type targeting via envelope engineering^19,24,53^, which could enable mtBE delivery to other tissues *in vivo*. Additionally, VLP technologies have been increasingly adopted by the gene editing field for genetic disease modeling and the generation of genetically engineered mouse models^46^; we expect that mtBE-VLPs will be readily compatible with these workflows and could enable the generation of additional mitochondrial disease models.

A single subretinal injection of mtBE-VLPs into adult mice enabled efficient mtDNA base editing in RPE cells. As an initial proof-of-concept demonstration, we showed *in vivo* installation of the LHON-disease-relevant *Nd4* m.G11185A mutation. Although there is a current scarcity of widely available mouse models of pathogenic mtDNA variants, more such models are actively being created using mtBEs^2,15,17^, which could enable further evaluation of potential therapeutic mtBE-VLP-mediated editing *in vivo*. Indeed, many pathogenic mtDNA variants have been implicated in ocular disorders^54^, highlighting the need for effective mtDNA editing and delivery strategies for ocular tissues. Notably, modified lentiviral particles packaging Cas9 mRNA, which share an envelope composition similar to that of mtBE-VLPs^33^, have entered clinical trials for the treatment of ocular disorders, with promising early results^55^. These developments, combined with our proof-of-concept *in vivo* data, suggest that continued optimization of mtBE-VLPs could enable the correction of pathogenic mtDNA mutations in ocular tissues.

Overall, our findings suggest the utility of mtBE-VLPs as a general platform for the study and potential treatment of genetic mitochondrial disorders. We anticipate that this platform will be broadly applicable to the growing family of mitochondrial base editors and to a wide range of pathogenic mtDNA point mutations.

---

**Extended Data Fig. 1.**
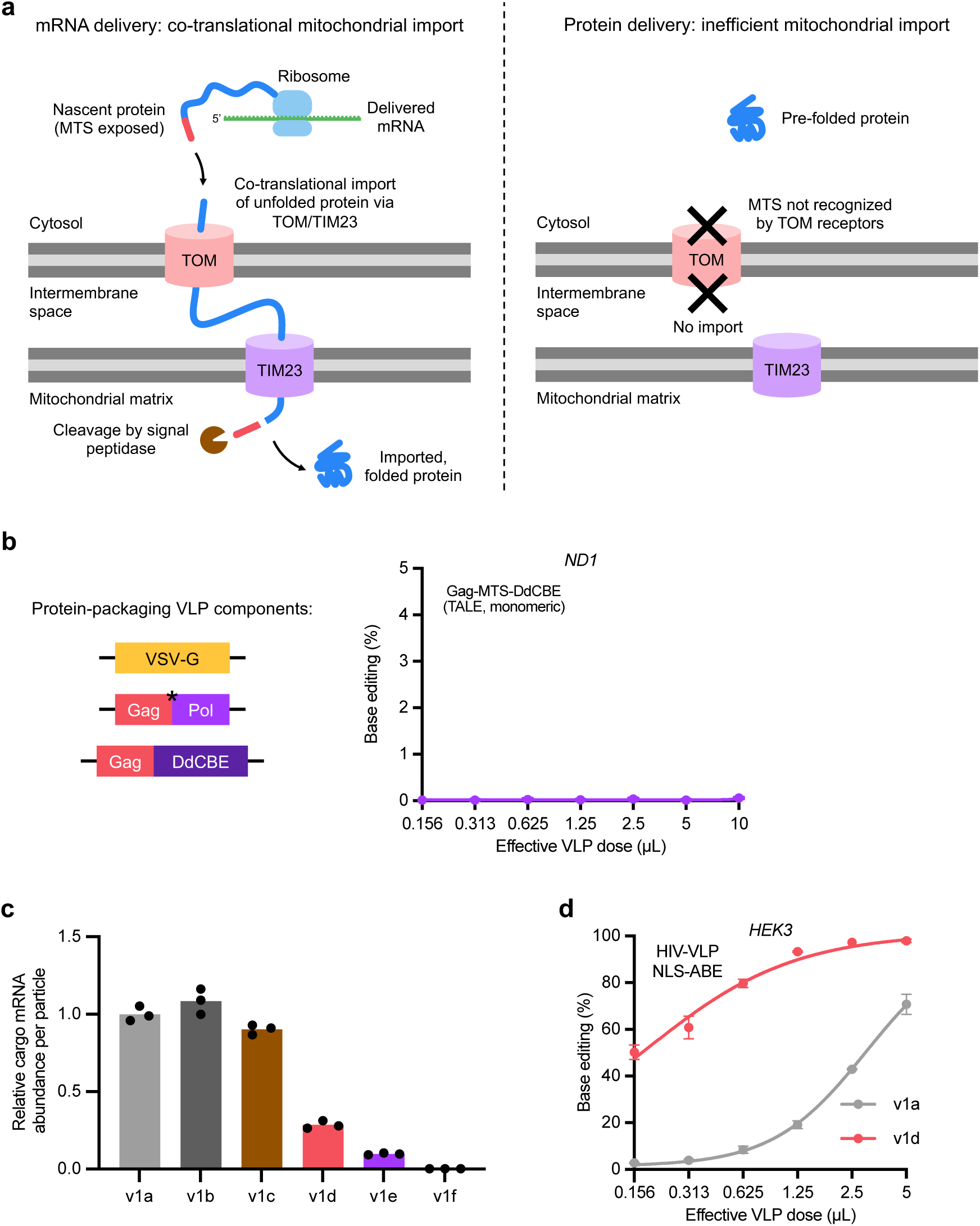
Comparison of protein-packaging and mRNA-packaging VLPs. **a,** Schematic diagram of co-translational mitochondrial import for delivery of mRNA versus protein. TOM, translocase of the outer mitochondrial membrane; TIM23, translocase of the inner mitochondrial membrane 23; MTS, mitochondrial targeting sequence. **b,** C•G-to-T•A base editing efficiencies at position G_10_ of the mitochondrial *ND1* locus in HEK293T cells by monomeric DdCBE protein-packaging VLPs. **c,** Relative abundance of cargo mRNA per particle across mtBE-VLPs produced using different Gag– Pol:Gag–MCP–Pol ratios, quantified by RT-qPCR and ELISA. Bars reflect the mean of n = 3 replicates, and dots represent individual replicate values. **d,** Comparison of nuclear ABE-packaging HIV-based VLPs produced using different Gag–Pol:Gag–MCP–Pol ratios. Adenine base editing efficiencies at position A_8_ of the *HEK3* target locus in HEK293T cells are shown. **b, d** Dots and error bars represent mean ± s.d. of n = 3 replicates. Data were fit to four-parameter logistic curves using nonlinear regression. Source data are provided as a Source Data file.

**Extended Data Fig. 2.**
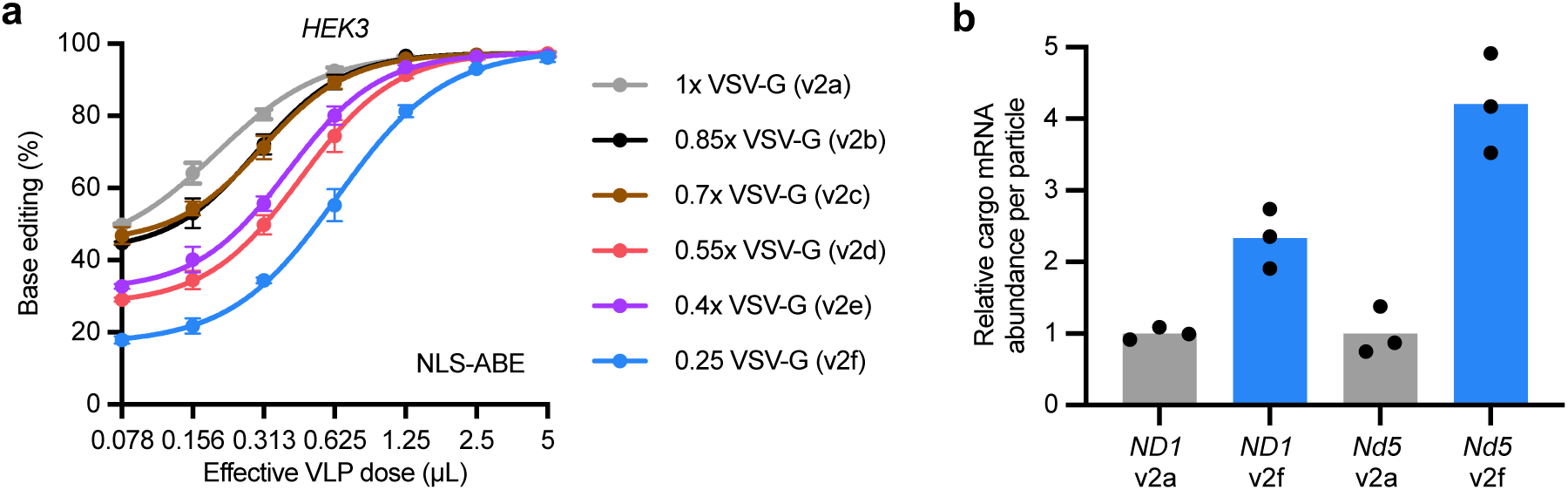
Reducing VSV-G expression negatively impacts the potency of nuclear ABE mRNA-packaging VLPs. **a,** Comparison of nuclear ABE mRNA-packaging VLPs produced using different VSV-G plasmid amounts. Adenine base editing efficiencies at position A_8_ of the *HEK3* locus in HEK293T cells are shown. Dots and error bars represent mean ± s.d. of n = 3 replicates. Data were fit to four-parameter logistic curves using nonlinear regression. **b,** Relative cargo mRNA abundance per particle across different mtBE-VLPs, quantified by RT-qPCR and ELISA. Bars reflect the mean of n = 3 replicates, and dots represent individual replicate values. Source data are provided as a Source Data file.

**Extended Data Fig. 3.**
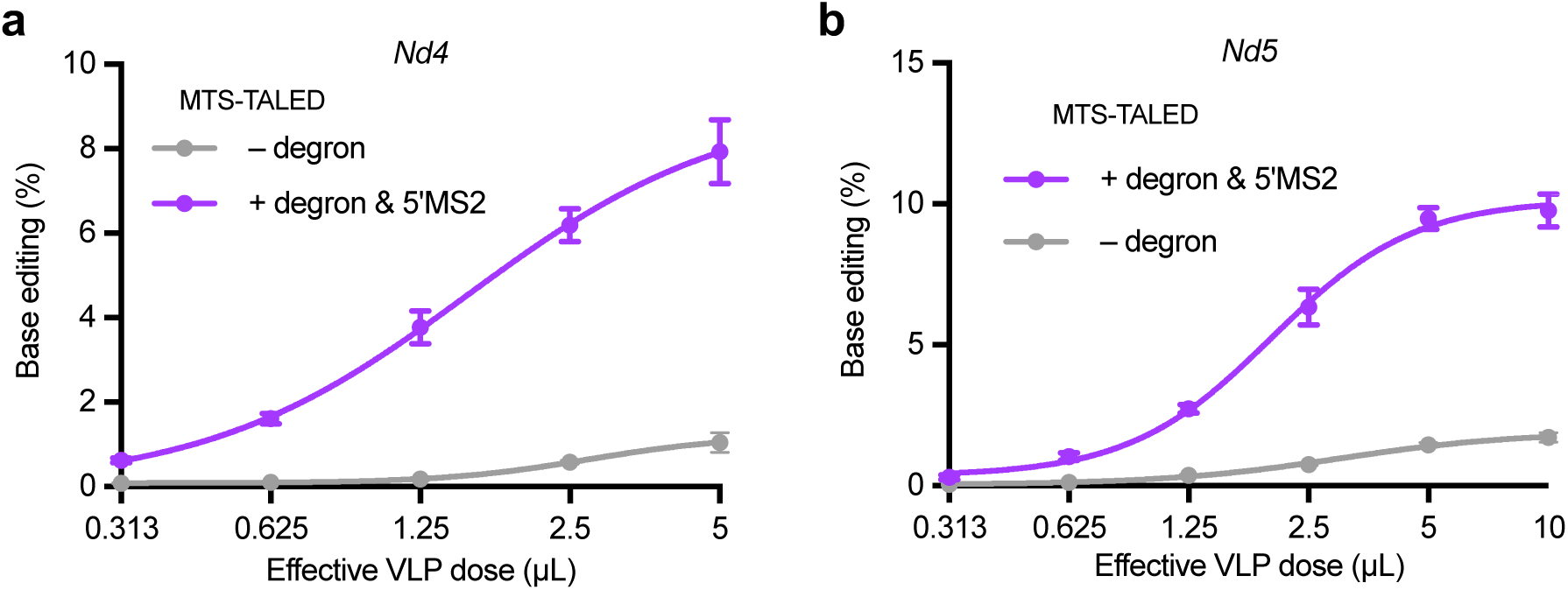
Comparison of TALED mtBE-VLPs at additional mtDNA target loci. A•T-to-G•C base editing efficiencies of unoptimized and degron/5’MS2-containing TALED mtBE-VLPs in mouse Neuro-2a cells at position A_11_ of the mitochondrial *Nd4* locus (**a**) and position A_12_ of the mitochondrial *Nd5* locus (**b**). Dots and error bars represent mean ± s.d. of n = 3 replicates. Data were fit to four-parameter logistic curves using nonlinear regression. Source data are provided as a Source Data file.

**Extended Data Fig. 4.**
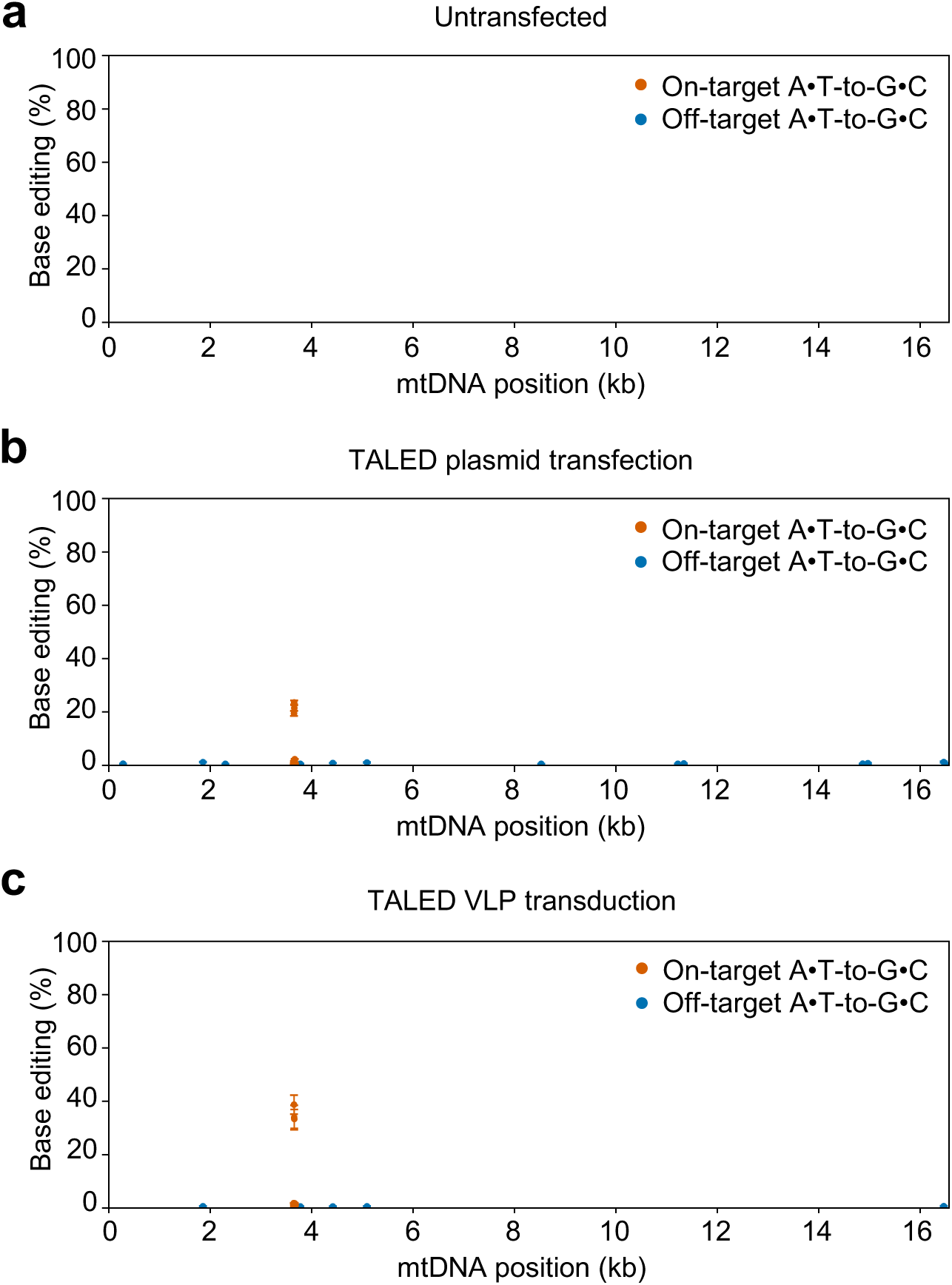
mtDNA-wide off-target editing generated by TALED mtBE-VLP transduction compared to plasmid transfection. **a,** Background levels of mtDNA-wide single-base A•T-to-G•C editing frequencies in untransfected HEK293T cells. **b,** mtDNA-wide single-base A•T-to-G•C editing frequencies in HEK293T cells following *ND1*-targeting TALED plasmid DNA transfection. **c,** mtDNA-wide single-base A•T-to-G•C editing frequencies in HEK293T cells following *ND1*-targeting TALED mtBE-VLP transduction. **a–c,** Each dot represents a single C•G base pair, and error bars represent mean ± s.d. of n = 3 replicates. Orange dots denote on-target editing; blue dots, off-target editing. Source data are provided as a Source Data file.

## Methods

### Cloning

Plasmids used in this study were cloned via Gibson cloning (NEBuilder® HiFi DNA Assembly; E2621X) or Golden Gate cloning. DNA was amplified via PCR using Phusion HotStart II polymerase (Thermo Fisher Scientific; F537L). To clone DdCBE and TALED constructs, Addgene plasmids #179680, #179685, #191510, and #191513 (DdCBE^8,11^, gifts from David R. Liu), as well as #183892, #183898, #169212, and #169213 (TALED^4,56^, gifts from Jin-Soo Kim). The other TALE-, and OpenABE-encoding sequences were synthesized by Integrated DNA Technologies based on sequences reported previously^15,42^. Mach1 (Thermo Fisher Scientific; C862003) chemically competent E. coli were used for plasmid propagation. Plasmids were extracted using QIAGEN Plasmid Plus Midi (12945) or Maxi (12963) Kits according to the manufacturer’s protocols.

### Cell culture

HEK293T cells (ATCC; CRL-3216), Gesicle Producer 293T cells (Takara; 632617), Neuro-2a cells (ATCC; CCL-131) were maintained in DMEM + GlutaMAX (Life Technologies; 10569044) supplemented with 10% (v/v) fetal bovine serum (Gibco; A5670701). Mouse embryonic fibroblasts (ScienCell; M7540-57) and GM10742 cells (Coriell Institute) were maintained in DMEM/F12 (ScienCell; 09421) and RPMI + GlutaMAX (Life Technologies; 61870036), respectively, supplemented with 10% (v/v) fetal bovine serum (Gibco; A5670701). Cells were cultured at 37 °C with 5% carbon dioxide and were confirmed to be negative for mycoplasma by testing with MycoAlert (Lonza; LT07-318).

### Production and purification of virus-like-particles

VLPs were produced as described previously^25,26^. Briefly, VLPs were produced by transient transfection of Gesicle 293T cells. Gesicle 293T cells were seeded in T-75 flasks at a density of 5 × 10^6^ cells per flask in 10mL of media. After 20–24 h, cells were transfected using the jetPRIME transfection reagent (Polyplus; 114-75) according to the manufacturer’s protocols. For producing v1a–v1f VLPs, a mixture of plasmids expressing VSV-G (760 ng), MMLV gag-MCP-pol and MMLV gag-pol (combined at 4270 ng per flask, mixed at 100:0, 80:20, 60:40, 40:60, 20:80, and 0:100 ratios for v1a through v1f, respectively), and cargo plasmid containing MS2 array (4270 ng) was co-transfected per T-75 flask. To assess the effect of VSVG plasmid amount on VLP production, the amount of VSV-G plasmid was reduced stepwise by 15% across six conditions, ranging from 760 ng to 190 ng. To maintain a constant total plasmid mass across all conditions, the amount of VSV-G reduced relative to the reference (760 ng) condition was redistributed equally between the gag-pol, gag-MCP-pol and cargo plasmids, preserving their original mass ratio. For production of VLPs containing degron-tagged cargo, dTAG-13 (Selleckchem; S9813) was added to the culture media 6 h post-transfection. Briefly, 40–48 h post-transfection, producer cell supernatant was harvested and centrifuged at 500 × g for 5 min to remove cell debris. The clarified eVLP-containing supernatant was then filtered through a 0.45-μm PVDF filter (Millipore; SE1M003M00). The filtered supernatant was subsequently concentrated by either PEG-it precipitation or ultracentrifugation. For PEG-it concentration, PEG-it Virus Precipitation Solution (SystemBiosciences; LV825A-1) was used according to the manufacturer’s protocol, and the resulting pellet was resuspended in Opti-MEM serum-free medium (Thermo Fisher Scientific; 31985070). For ultracentrifugation, the supernatant was layered over a 20% (w/v) sucrose/PBS cushion and centrifuged at 26,500 rpm (86,242 × g) for 2 h at 4°C using an SW32 Ti rotor in an Optima XE ultracentrifuge (Beckman Coulter). Following ultracentrifugation, VLP pellets were resuspended in cold PBS, pH 7.4 (Gibco; 10010023). For subretinal injections, the resuspended VLPs were additionally centrifuged at 9000 × g for 5 min to remove debris. VLPs were then frozen at a rate of −1°C/min and stored at −80°C. VLPs were thawed on ice immediately prior to use.

### Generation of gRNA-expressing HEK293T cells

The *HEK3*-targeting gRNA sequence was cloned into a lentiviral transfer vector carrying a puromycin resistance gene. To produce lentivirus, HEK293T cells were seeded in a T-25 flask at a density of 2 × 10^6^ cells per flask. The following day, cells were co-transfected with 2 μg VSV-G envelope plasmid, 3 μg psPAX2 packaging plasmid and 3 μg transfer vector encoding the *HEK3* gRNA, using 23 μL FuGENE in 0.5 mL Opti-MEM. Two days after transfection, the culture medium was collected, centrifuged (500g, 5 min) to remove cell debris, and filtered through a 0.45-μm PVDF membrane. HEK293T cells were then transduced with the lentivirus-containing medium. One day after transduction, untransduced cells were eliminated by culturing in medium supplemented with 2 μg/mL puromycin for 5 days. The *HEK3* gRNA sequence is listed in Supplementary Table 1.

### VLP transduction in cell culture and genomic DNA isolation

Cells were plated for transduction in 96-well plates (Corning) at a density of 16,000 cells per well. After 20–24 h, VLPs were added directly to the culture media in each well. 72 h post-transduction, cellular genomic DNA was isolated as previously reported^25^. Briefly, media was aspirated from each well, and 60 µL of lysis buffer (10 mM Tris-HCl pH 8.0, 0.05% SDS, 25 µg mL−1 Proteinase K (Thermo Fisher Scientific; EO0492)) was added to each well. Lysis was achieved by incubation at 37 °C for 1–1.5 h followed by heat inactivation at 80 °C for 30 min.

### High-throughput sequencing of genomic DNA

Genomic DNA was isolated as described above. 1 µL of the isolated DNA was used as input for the first of two PCR reactions. Genomic loci were amplified in PCR1 using Phusion U polymerase (Thermo Fisher Scientific; F562L). PCR1 primers are listed in Supplementary Table 2 under the HTS_forward and HTS_reverse columns. PCR1 was performed as follows: 95 °C for 3 min; 16 cycles of 95 °C for 15 s, 56 °C for 20 s, and 72 °C for 30 s; 72 °C for 1 min. In all, 1 μL of PCR1 was used as an input for PCR2 to install Illumina barcodes. PCR2 was conducted for nine cycles of amplification using Phusion HotStart II polymerase (Thermo Fisher Scientific; F537L). Following PCR2, samples were pooled and gel purified in a 1% agarose gel using a QIAquick Gel Extraction Kit (Qiagen; 28704). Library concentration was quantified using the QubitTM dsDNA High-Sensitivity Assay Kit (Thermo Fisher Scientific; Q33230). Samples were sequenced on Illumina MiSeq or Miseq i100 instruments using Illumina MiSeq Reagent Kit v2 (Illumina; MS-102-2002) or MiSeq i100 Series 5M Reagent Kit (Illumina; 20126565).

### High-throughput sequencing data analysis

Sequencing reads were demultiplexed using the MiSeq Reporter software (v2.6) (Illumina) and were analyzed using CRISPResso2 (v2.3.2)^57^ as previously described^25^. Batch analysis mode (one batch for each unique amplicon analyzed) was used in all cases. Reads were filtered by minimum average quality score (Q > 30) prior to analysis. For analysis of base editing efficiencies, the following quantification window parameters were used: -w 20 -wc -10. Even when no sgRNA was used in the experiment (i.e. for TALE or ZF-based mtBEs), a protospacer sequence was used after the -g flag to enable quantitation. Base editing efficiencies are reported as the percentage of sequencing reads containing a given base conversion at a specific position. Target base positions within a given amplicon are numbered relative to the first position of the protospacer sequence used for CRISPResso2 analysis, listed in Supplementary Table 2. Prism 11 (GraphPad) was used to generate dot plots and bar plots. Amplicon sequences are provided in Supplementary Table 2.

### *In vitro* transcription of spike-in RNA controls

Firefly luciferase (FLuc) RNA was prepared by in vitro transcription using the HiScribe® T7 Quick High Yield RNA Synthesis Kit (New England Biolabs; #E2050S), following the manufacturer’s protocol. The FLuc control template DNA, containing the firefly luciferase gene under the control of the T7 promoter, was provided in the kit and used as the template for in vitro transcription. The resulting FLuc RNA was used for spike-in RNA for RT-qPCR.

### VLP RNA extraction and RT-qPCR

Total RNA was extracted from VLPs using the QIAamp Viral RNA Mini Kit (Qiagen; 52904), with Firefly luciferase (FLuc) spike-in RNA included during extraction. Extracted RNA was reverse transcribed using SuperScript™ III First-Strand Synthesis SuperMix (Thermo Fisher Scientific; 18080400) with random hexamer primers, according to the manufacturer’s protocol. qPCR analysis of the resulting cDNA was performed on a QuantStudio™ 3 Real-Time PCR System (Thermo Fisher Scientific) using SYBR Green dye (Lonza; 50512). Relative cargo mRNA abundance was calculated using the ΔΔCt method, with FLuc used as the normalization control. qPCR primers are listed in Supplementary Table 1.

### Quantification of VLP titer

To quantify particle number, the concentration of MLV p30 protein in purified VLPs was determined using the MuLV Core Antigen ELISA Kit (Cell Biolabs; VPK-156), according to the manufacturer’s protocol. To calculate VLP titer in particles/µL, the p30 concentration in ng/mL determined by ELISA was multiplied by a factor 2030.57, which accounts for molarity conversions, assumes that 20% of p30 molecules in solution are associated with particles, and assumes a copy number of 1800 molecules of p30 per VLP, as previously described^58^.

### Whole mitochondrial genome sequencing

For whole mitochondrial genome sequencing, genomic DNA was initially isolated as described above. mtDNA was then amplified by PCR using Phusion U polymerase (Thermo Fisher Scientific; F562L) and two sets of slightly overlapping primers to reduce primer bias. Each primer pair amplified almost 50% of the mtDNA. Primer sequences are provided in Supplementary Table 1. The PCR products were then purified with a QIAquick PCR purification kit (Qiagen; 28104). Finally, we utilized an Illumina DNA Prep kit (Illumina; 20060060) with Nextera DNA CD Indexes to create an NGS library from the purified PCR products (Illumina). The libraries were then pooled and transferred onto a MiSeq sequencer (Illumina).

### Analysis of mitochondrial genome-wide off-target editing

To analyze NGS data from whole mitochondrial genome sequencing, we adopted a method previously used to investigate off-target effects in mitochondrial genomes^4,5,7,8^. Initially, we aligned the Fastq sequences to the mitochondrial (MT) sequence extracted from the GRCh38 (release v102) reference genome using BWA (v.0.7.17), and then created BAM files with SAMtools (v.1.9) by fixing read pairing information and flags. Subsequently, we used the REDItoolDenovo.py script from REDItools (v.1.2.1) to identify all cytosines, guanines, adenines and thymines in the mitochondrial genome exhibiting conversion rates > 0.1%. Positions with conversion rates ≥ 5% in both treated and untreated samples were identified as SNVs in the cell lines and removed, along with the construct’s on-target sites. The remaining positions were considered off-target sites, and the average off-target editing frequency was calculated as the total number of edited C-to-T and G-to-A nucleotides for DdCBE, or A-to-G and T-to-C nucleotides for TALED, divided by the total coverage across all off-target sites. Mitochondrial genome-wide graphs were constructed by plotting the conversion rates at on-target and off-target sites with an editing frequency > 1% for DdCBE and > 0.5% for TALED across the entire mitochondrial genome. The custom Python scripts used for this analysis are available at “https://github.com/hjang-wi/mt-offtarget”.

### Producer-cell protein extraction and Western blot

For protein extraction from producer cells, 24 h after transfection, cells were trypsinized and 1 × 10^6^ cells were collected by centrifugation at 500× g for 5 min. Cell pellets were washed once with 1 mL of cold PBS and lysed using RIPA buffer (Thermo Fisher Scientific; 89901) supplemented with cOmpleteTM protease inhibitor cocktail (Roche Diagnostics; 11836153001) according to the manufacturer’s protocols. Cells were lysed at 4 °C for 30 min with gentle agitation. The lysate was clarified by centrifugation at 10,000× g for 20 min at 4 °C, and the clarified supernatant was combined with 4× Laemmli Sample Buffer (Bio-Rad; 1610747) and heated at 95 °C for 10 min. Protein extracts were subsequently separated by electrophoresis at 150 V for 45 min on a NuPAGE 3–8% Tris-Acetate gel (Thermo Fisher Scientific; EA0378BOX) in 1× NuPAGE Tris-Acetate SDS running buffer (Thermo Fisher Scientific; LA0041). Transfer to a PVDF membrane was performed using an iBlot 2 Gel Transfer Device (Thermo Fisher Scientific) at 20 V for 7 min. The membrane was blocked for 1 h at room temperature with rocking in blocking buffer: 2% bovine serum albumin (BSA) and 0.1% Tween-20 in TBS (150 mM NaCl and 50 mM Tris-HCl). After blocking, the membrane was incubated overnight at 4 °C with rabbit anti-HA (Cell Signaling Technology; 3724, 1:1000 dilution) and mouse anti-actin (Cell Signaling Technology; 3700, 1:1000 dilution). The membrane was washed three times with 1×TBST (150 mM NaCl, 0.1% Tween-20, and 50 mM Tris-HCl) for 10 min each time at room temperature, then incubated with goat anti-mouse secondary antibody (LI-COR IRDye 680RD; 926-68070, 1:10,000 dilution in blocking buffer) and goat anti-rabbit secondary antibody (LI-COR IRDye 800CW; 926-32211, 1:10,000 dilution in blocking buffer) for 1 h at room temperature with rocking. The membrane was washed as before and imaged using an Odyssey Imaging System (LI-COR). Protein abundances were quantified by densitometry using ImageJ (v2.14.0). protein abundances were normalized to the actin abundance within each sample, and comparisons across samples were performed using these actin-normalized values.

### TMRE assay of producer cells

Gesicle producer cells were plated at 0.5 × 10⁶ cells per 35-mm glass-bottom dish and transfected as described above, with all plasmid and transfection reagent amounts scaled down proportionally to the reduced cell number. At 24 h post-transfection, mitochondrial membrane potential was assessed using the TMRE-Mitochondrial Membrane Potential Assay Kit (APExBIO; K2233) according to the manufacturer’s protocol, with CCCP-treated cells and untransfected cells serving as positive and negative controls, respectively. Cells were stained with TMRE working solution for 60 min at 37 °C protected from light, washed twice with pre-warmed medium, and imaged using a Dragonfly 505 spinning-disk confocal microscope (Andor Technologies) equipped with a piezo Z-stage (ASI) and an iXon Ultra 888 EMCCD camera, at an excitation/emission wavelength of 550/575 nm.

### Cell viability assays

Cell viability assays were performed using CellTiter-Glo® (Promega; G7570) at 24 h after plasmid transfection, following the manufacturer’s protocol. Luminescent signal was quantified using a SpectraMax® iD5 microplate reader.

### RNA sequencing and data analysis

At 24 h post-transfection, approximately 1x10^5^ cells were collected by centrifugation, resuspended in 50 μL Zymo DNA/RNA Shield™ (Zymo Research; R1100-50), and submitted for Plasmidsaurus RNA-seq. Briefly, sequencing libraries were prepared using an Illumina-based 3’ end counting approach. mRNA was converted into complementary DNA (cDNA) via reverse transcription using oligo(dT) primer and second-strand synthesis, followed by tagmentation, library indexing, and sequencing. For RNA A-to-G off-target editing analysis, mitochondrial adenosine positions were called using REDItools (v1.2.1) and A-to-G editing frequencies were computed directly from the BaseCount field. Sites detected in any untreated control replicate were considered germline SNPs and excluded from all samples. Sites with an A-to-G editing frequency ≥0.5% were considered as edited. The custom Python scripts used for RNA sequencing analysis are available at “https://github.com/hjang-wi/mt-offtarget”.

### Spinoculation of LHON patient-derived LCLs

LHON patient-derived LCL cells (Coriell; GM10742) were transduced with VLPs via spinoculation. Cells were collected by centrifugation, counted, and seeded at 1 × 10⁴ cells per well in U-bottom 96-well plates. Polybrene was added to a final concentration of 4 µg/mL, followed by addition of VLPs. Plates were then centrifuged at 800 × g for 60 min at 32°C using a swinging-bucket rotor. Following spinoculation, plates were gently tapped to resuspend the cell pellet and incubated at 37°C with 5% CO₂. Genomic DNA was extracted 3 days post-transduction.

### Animals

C57BL/6J (“WT,” JAX 000664) mice were purchased from the Jackson Laboratory (Bar Harbor, ME). Mice were housed in the vivarium at University of California, Irvine, and maintained on a normal mouse-chow diet and a 12 h/12 h light/dark cycle. All animal procedures were approved by the Institutional Animal Care and Use Committee of the University of California, Irvine; and were conducted in accordance with the guidelines of the NIH for the care and use of laboratory animals, and with the Statement for the Use of Animals in Ophthalmic and Visual Research by the Association for Research in Vision and Ophthalmology.

### Subretinal injection

4-week-old mice were used for all experiments, with equal numbers of male and female mice randomized across experimental groups. Before subretinal injection, the mouse eyes were treated with a topical administration of 1% tropicamide ophthalmic solution (Akorn; 17478-102-12) and 10% phenylephrine (Valeant; 42702-0103-05) to effect pupil dilation. After adequate dilation, mice were anesthetized by intraperitoneal injection of a cocktail consisting of 20 mg mL−1 ketamine and 1.75 mg mL−1 xylazine in PBS at a dose of 0.1 mL per 20 g body weight. Subretinal injections were performed under an ophthalmic surgical microscope (Zeiss). Corneal hydration was achieved with topical application of Eye Lube Plus Lubricating Gel (OptixCare). An incision was made through the cornea adjacent to the limbus at the nasal side using a 27-gauge needle. A 34-gauge blunt-end needle (World Precision Instruments; NF34BL) connected to an RPE-KIT (World Precision Instruments; RPE-KIT) by SilFlex tubing (World Precision Instruments; SILFLEX-2) was inserted through the corneal incision while avoiding the lens, and advanced through the retina. Each mouse was injected with 1.5 μl of PBS or VLPs (approximately 6.7 × 10^10^ particles). After injections, the gel was reapplied, anesthesia reversal was achieved with intraperitoneal atipamezole (2.5 mg kg^−1^; MWI Animal Health; 032800), and mice were allowed to recover on a heat pad.

### RPE dissociation and analysis of RPE genomic DNA

Seven days after injection, mouse eyes were dissected under a light microscope to separate the posterior eyecup (containing RPE, choroid, and sclera) from the retina and anterior segment. Each posterior eyecup was immediately immersed in PBS. RPE cells were detached in PBS from the posterior eyecup (choroid and scleral cells) by gentle pipetting, followed by removal of the remaining posterior eyecup. Genomic DNA was purified using the DNeasy Blood & Tissue Kit (Qiagen; 69504), according to the manufacturer’s protocol. 100 ng of purified genomic DNA was amplified for high-throughput sequencing using mitochondrial gene-specific primers as described above, using 16 cycles of PCR1 amplification.

### Statistics and reproducibility

Data are presented as mean and standard deviation (s.d.). Statistical analysis was performed using Prism 11 (GraphPad). Sample sizes are described in the figure legends.

### Data availability

The high-throughput sequencing data generated during this study is being deposited at the NCBI Sequence Read Archive database under PRJNA1522533. Source Data is provided with this paper. The protein sequences for all mtBE constructs used in this study are listed in Supplementary Table 3. Key plasmids from this work will be deposited to Addgene for distribution.

### Code availability

The code used for analyzing base editing efficiencies are available at https://github.com/pinellolab/CRISPResso2.

## Acknowledgements

This work was supported in part by the Whitehead Institute and Whitehead Fellows Program (A.R.); NIH Director’s Early Independence Award DP5OD037342 (A.R.); McGuire Family Foundation (A.R.); Owens Family Foundation (A.R.); Valhalla Foundation (A.R.); NIH R01EY009339 (K.P.); and NIH R01EY030873 (K.P.). H.J. acknowledges support from the National Research Foundation of Korea. C.R.M is a Beckman Scholar and acknowledges support for the research described in this study from the Arnold and Mabel Beckman Foundation. We also acknowledge support to the Department of Ophthalmology Gavin Herbert Eye Institute at the University of California, Irvine from an unrestricted Research to Prevent Blindness award, from NIH core grant P30EY034070, and from a University of California, Irvine School of Medicine Dean’s Office grant. We thank S.-I. Cho for assistance with TALE design; Y. Chang and A. Jain for assistance with confocal imaging; D. Liu and the Whitehead Genome Technology Core for generous access to Illumina sequencing instrumentation; and the Jain, Reddien, Weissman, and Young Labs for generous access to equipment.

## Author contributions

H.J. and A.R. conceived and designed the study. H.J. developed mRNA-packaging VLP architectures, performed all cell culture experiments, and analyzed data. C.R.M. and K.P. designed and performed the mouse subretinal injection experiments. H.J. and A.R. wrote the manuscript with input from all authors.

